# Development of a replication competent murine norovirus reporter system

**DOI:** 10.1101/2025.01.15.633260

**Authors:** Mikayla C. Olson, Robert C. Orchard

**Author notes:** Correspondence to (R.C.O.).

## Abstract

Caliciviruses are significant agricultural and human pathogens that are poorly understood due to the dearth of molecular tools, including reporter systems. We report the development of a stable, faithful, and robust luciferase-based reporter system for a model calicivirus, murine norovirus (MNoV). Genetic insertion of a HiBiT tag, an 11 amino acid fragment of nanolucifersase, at the junction of the nonstructural proteins NS4 and NS5 yields infectious virus. The resultant MNoV-HiBiT produces robust signal that is detected early in infection and occurs only in cells susceptible to MNoV infection. As proof of principle, we used this tool to characterize two unappreciated host directed anti-MNoV compounds. The use of the MNoV-HiBiT virus enables new mechanistic studies by a rapid and quantitative means of measuring MNoV replication. Furthermore, the HiBiT insertion strategy we describe may be useful for the generation of other calicivirus reporters.

**Author Summary:** Colloquially known as “the stomach bug,” human norovirus is a leading cause of foodborne illness worldwide. Despite its prevalence, there are no approved therapeutics or vaccines against norovirus owing to the challenges in cultivation of the virus and a lack of a small animal model of human norovirus. Murine Norovirus (MNoV) has emerged as a tractable model due to its similarity to human norovirus and the ability to grow the virus in laboratory strains of mice and immortalized cells in vitro. Reporter viruses, which include enzymatic or fluorescent reporter genes in the viral genome, are invaluable tools that expedite viral research. Unfortunately, there are no MNoV reporter systems available. Here we developed a novel reporter MNoV using a complimentary nanoluciferase. Specifically, we inserted a HiBiT tag into the MNoV genome that is functionalized into a full nanoluciferase protein and emits light in the presence of LgBiT. We demonstrated that MNoV-HiBiT is a faithful and sensitive reporter of viral replication across MNoV strains and cell types. We used the reporter virus to screen novel anti-viral compounds, demonstrating that the MNoV-HiBiT system is a powerful tool to enhance norovirus research.

## Introduction

Human Norovirus (HNoV), a positive-sense RNA virus of the *Calicivirdae* family, is the leading cause of gastroenteritis worldwide [1]. Currently, there is no approved vaccine or therapeutic for HNoV, largely due to our limited understanding of calicivirus biology [1]. For many viral systems, the development of reporter viruses has greatly enhanced our understanding of the viral life cycle and accelerated the development of therapeutics and vaccines. Reporter systems for caliciviruses are lacking. Unlike HNoV, murine norovirus (MNoV) replicates in immortalized tissue culture cell lines and can infect laboratory strains of mice [2, 3]. Thus, MNoV has emerged as the premier calicivirus model system. Despite a robust reverse genetic system, the incorporation of fluorescent or enzymatic reporters into the MNoV genome has not been successful. Recently, a single cycle MNoV reporter system was developed through a trans complementation approach [4]. However, a replication competent reporter system for caliciviruses has yet to be achieved and would advance research progress.

## Results

We were inspired by the previously generated T-cell epitope viruses where the small peptide SIINFEKL or the GP33 epitope is inserted between MNoV nonstructural proteins 4 and 5 (NS4 and NS5) with a duplicated protease cleavage site [5, 6]. To generate a replication competent reporter system, we inserted the small HiBiT peptide sequence at the identical site into the genome of MNoV^CW3^ (herein called MNoV^CW3^-HiBiT; **Figure 1A**). The eleven amino acid HiBiT tag can be functionalized into nanoluciferase by the addition of LgBiT in cells or in the luciferase lysis buffer [7]. The latter enables the use of HiBiT system in any cell type without the need to modify the cells. The HiBiT epitope is detected on NS4-5 polyprotein precursors and to a small extent on processed NS4. (**Figure 1B**). The molecular clone of MNoV^CW3^-HiBiT produced infectious virus. Insertion of the HiBiT sequence significantly diminished MNoV^CW3^-HiBiT replication but still enabled exponential growth of the virus compared to parental MNoV^CW3^ (**Figure 1C**). To assess the functionality and specificity of MNoV^CW3^-HiBiT in vitro, we performed a time course luciferase assay using either wildtype BV2 cells or BV2ΔCD300lf cells, which lack the receptor necessary for MNoV infection [8]. Cells were inoculated with either the parental MNoV^CW3^ or MNoV^CW3^-HiBiT virus and lysed after a single cycle of infection (12 hours) or multiple cycles of infection (24 or 48 hours). We then measured the luminescence from the cell lysate by supplying the LgBiT via the lysis buffer *in trans*. Infection of wild-type BV2 cells with MNoV^CW3^-HiBiT led to exponential increase in light production by luciferase with nearly a 1000-fold difference in RLU at 48 hours post-infection (**Figure 1D**). This signal was specific to the HiBiT virus and required entry of MNoV (**Figure 1D**). To further assess the sensitivity of the reporter virus, we conducted another time course assay with a high multiplicity of infection (MOI) and timepoints at 1-hour increments for 12 hours. Statistical differences in luminescence for MNoV^CW3^-HiBiT in wildtype BV2 cells compared to controls were detected as early as at 8 hours post-infection (**Figure 1E**). Taken together, these data demonstrate that the MNoV^CW3^-HiBiT virus is specific in monitoring viral replication by luminescent output.

**Figure 1:**
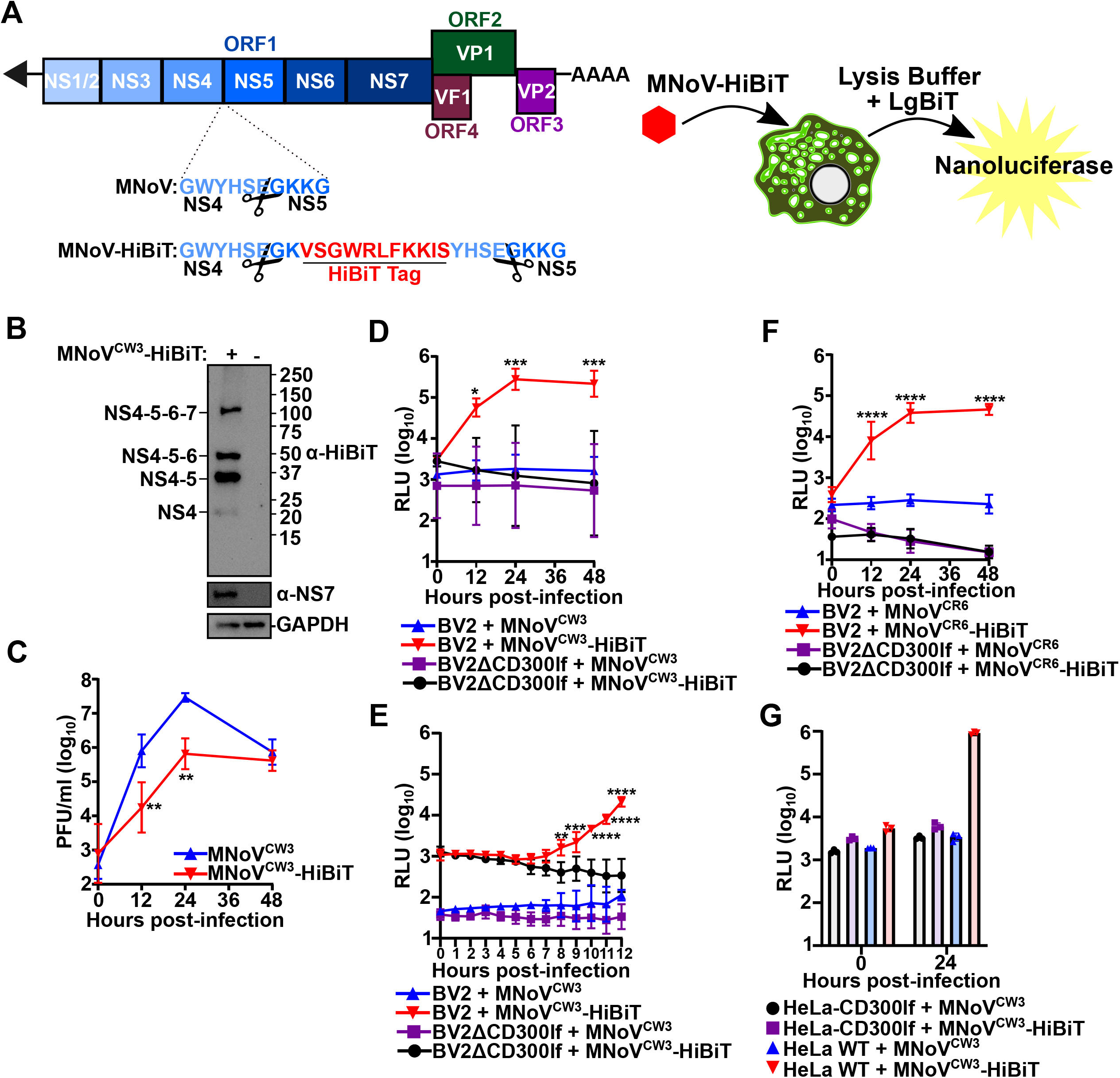
Development and Characterization of MNoV-HiBiT Viruses. **A)** (Left) Cartoon illustration of the MNoV genome organization with a focus on the NS4-NS5 cleavage site and the introduction of the HiBiT tag (red). Scissors indicate the MNoV protease cleavage site. (Right) Cartoon illustration of the utility of the MNoV-HiBiT virus and the complementation with LgBiT supplied in lysis buffer to form a functional nanoluciferase. **B)** Representative western blot of BV2 cells either uninfected or infected with MNoV^CW3^-HiBiT for 12 hours and probed for indicated antibodies. Predicted molecular weight of MNoV nonstructural proteins are indicated. **C)** BV2 cells were challenged with MNoV^CW3^ or MNoV^CW3^-HiBiT at a multiplicity of infection (MOI) of 0.05. Viral production was enumerated using plaque assays (PFU; plaque forming units) at the indicated time points. **D-F)** BV2 or BV2ΔCD300lf cells were challenged with MNoV^CW3^ or MNoV^CW3^-HiBiT (**D-E)** or MNoV^CR6^ or MNoV^CR6^-HiBiT (**F)** at a multiplicity of infection (MOI) of 5.0. At the indicated time points, infection was quantified using Nano-Glo HiBiT Lytic Detection System (RLU; relative luminescent units). **G)** HeLa or HeLa-CD300lf cells were challenged with MNoV^CW3^ or MNoV^CW3^-HiBiT at an MOI of 1.0 and at the indicated time points, infection was quantified using Nano-Glo HiBiT Lytic Detection System. All data are shown as mean ± S.D. from three independent experiments and analyzed by one way ANOVA with Tukey’s multiple comparison test. Statistical significance is annotated as follows: ns not significant, * P < 0.05, ** P< 0.01, *** P< 0.001, **** P<0.0001.

MNoV strains exhibit different infection outcomes after oral inoculation [9]. For example, MNoV^CW3^ causes an acute infection that spreads to extraintestinal tissues and is lethal in STAT1 deficient animals. In contrast, MNoV^CR6^ establishes a persistent infection in the gastrointestinal tract and does not spread or cause lethality in immunodeficient animals [9, 10]. We set out to determine if our reporter system works across MNoV viral strains and target cells *in vitro*. First, we generated an MNoV^CR6^-HiBiT virus analogous to the one generated with MNoV^CW3^ described above. MNoV^CR6^-HiBiT had exponential growth of RLUs both at single and multicycle timepoints, indicating that this tagging strategy can be used for multiple MNoV strains (**Figure 1F**). Additionally, we validated that MNoV^CW3^-HiBiT is infectious and luminescent in human cells when CD300lf is ectopically expressed. Wildtype HeLa cells, which lack the MNoV receptor, or HeLa-CD300lf cells were inoculated with MNoV^CW3^ or MNoV^CW3^-HiBiT. The HeLa-CD300lf cells infected with MNoV^CW3^-HiBiT produced 1000-fold more RLUs than controls (**Figure 1G**). Taken together these data provide evidence that the MNoV-HiBiT system is robust across multiple MNoV strains and cellular systems.

Reporter viruses enable the assessment and screening of genetic or chemical perturbations at a scale that is challenging with traditional viral titer measurements. To demonstrate the utility of MNoV-HiBiT to identify new chemical inhibitors of norovirus replication, we mined a previously conducted CRISPR/Cas9 loss of function screen for MNoV dependency factors that have small molecule inhibitors [11]. We focused on Casein Kinase II (CKII) and CERT1 (also known as COL4A3BP) which have well characterized inhibitors: GO289 and HPA-12 respectively. Importantly, these molecules and their respective targets have not been validated to have anti-norovirus activity. We compared the antiviral activity of these compounds to the nucleoside analog 2′-C-methylcytidine (2CMC), which has known anti-norovirus activity [12, 13]. 2CMC inhibited MNoV^CW3^-HiBiT infection of BV2 cells with an IC_50_ of 0.68 µM, a value similar to that described previously for MNoV (**Figure 2A**) [12]. Treatment of BV2 cells with the CKII inhibitor GO289 (IC_50_ of 0.26 µM) and CERT inhibitor HPA-12 (IC_50_ of 8.19 µM) strongly reduced viral replication as measured by HiBiT-mediated luminescence, although HPA-12 displayed more cell toxicity (**Figure 2B and 2C**). Overall, these data demonstrate the utility of the MNoV-HiBiT system for identification and characterization of small molecule inhibitors of MNoV.

**Figure 2:**
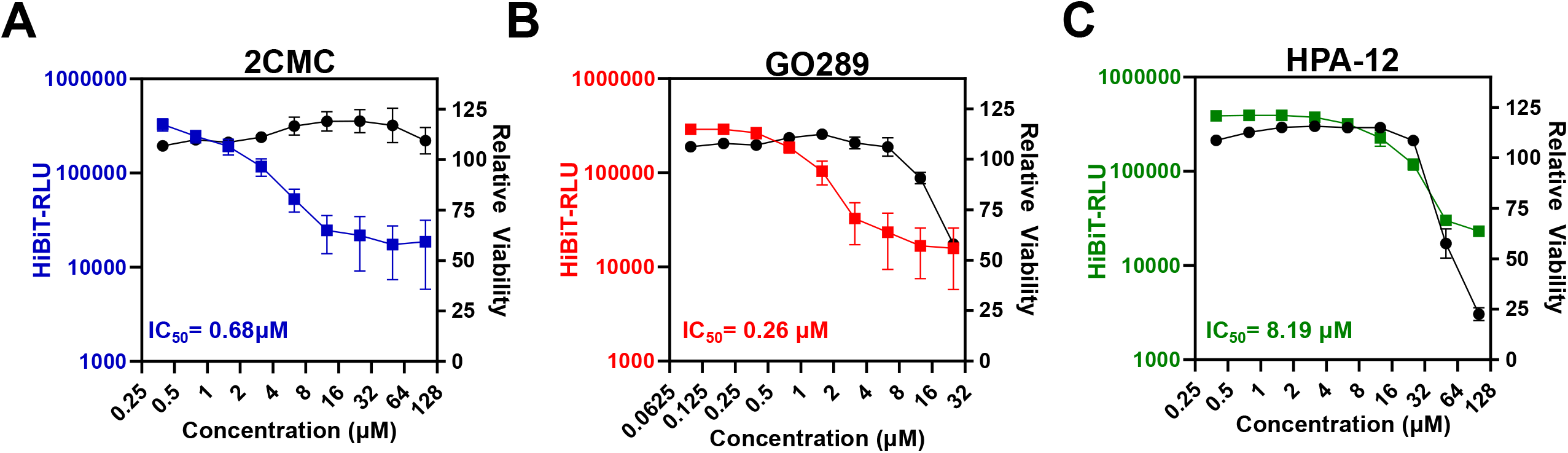
Identification of Anti-MNoV Activity of GO289 and HPA-12 Using MNoV-HiBiT System. **A-C)** Dose response curves of BV2 cells treated with indicated doses of 2CMC **(A)** GO289 **(B)**, or HPA-12 **(C)** infected with MNoV^CW3^-HiBiT (left axis) or uninfected and measured for cell viability using CellTiterGlo (right axis). IC_50_ are listed in bottom left corner. All data are shown as mean ± S.D. from three independent experiments.

## Discussion

Norovirus imposes a global health burden every year [14]. As such, novel tools to study viral replication and discover new therapeutics are necessary. Here, we have developed a replication competent reporter MNoV. MNoV-HiBiT confers a broad range of utility as the LgBiT can be added in the lysis buffer, eliminating the need for specialty cell lines. However, there are a few limitations of our current reporter system. First, the MNoV-HiBiT virus is significantly attenuated and thus caution in propagating this virus is necessary to avoid losing the inserted sequence. However, MNoV-HiBiT still has robust infection as evidenced by multiple MNoV strains and different cellular systems that we successfully interrogated. Second, the dual cleavage sites present uncertainty of the extent to which NS4 and NS5 are labeled compared to unlabeled, as we do not have specific antibodies targeting these nonstructural proteins. Also, we did not explore whether other insertion sites or strategies in the MNoV genome would have equal or superior sensitivity to the current design. The NS4-NS5 cleavage site is poorly processed across norovirus genogroups, including human norovirus [15]. Thus, it is possible that this insertion strategy may be broadly applicable to related caliciviruses. Taken together with the high dynamic range of luminescent output, MNoV-HiBiT can be useful across a multitude of applications. Elucidation of viral mechanisms, large scale screens, and antiviral drug validation, as shown here, are among the many opportunities for which MNoV-HiBiT can be instrumental.

## Materials and Methods

### Cell Culture

293T (ATCC), Hela (ATCC) and BV2 cells (Kind gift of Dr. Skip Virgin, Washington University) were cultured in Dulbecco’s Modified Eagle Medium (DMEM) with 5% fetal bovine serum (FBS). 10 µg/mL of blasticidin (Thermo Fisher) was added as appropriate. BV2ΔCD300lf and HeLa-CD300lf cells have been described previously [8]. Cell lines were tested regularly and verified to be free of mycoplasma contamination.

### Construction of MNoV-HiBiT

Plasmids encoding MNoV-HiBiT were generated in MNoV^CW3^ (Gen bank accession no. EF014462.1) or MNoV^CR6^ (Gen bank accession no. JQ237823) appropriately. The HiBiT tag with a cleavage site sequence on both sides was inserted between the coding regions for NS4 and NS5. DNA constructs were all verified by sequencing. MNoV-HiBiT plasmids were transfected into 293Ts and amplified on BV2s to generate infectious viruses as described previously [8].

### MNoV Assays

MNoV growth curve assays were conducted by seeding 5 × 10^4^ BV2 cells in 96-well plates and infecting with virus in suspension. Plates were frozen at -80C at the appropriate timepoints before performing plaque assay as previously described [8]. Briefly, 2.5 × 10^4^ BV2 cells were seeded per well of 24-well plate. Media was removed the following day, and 10-fold serial dilutions of lysate were added to each well, incubated with gentle rocking for 1 hour, and then removed. Cells were overlayed with complete media containing 1% methylcellulose. Plates were incubated for 3 days before staining with crystal violet solution.

### Luciferase Assay

Luminescent activity was measured with a Synergy LX multi-mode reader (Biotek) using a Nano-Glo HiBiT Lytic Detection System (Promega N3040) following the manufacturer’s protocols. 5 × 10^4^ BV2 or ΔCD300lf BV2 cells were seeded in 96-well plates and the next day inoculated with virus and frozen at -80°C at the indicated time point. For Hela and CD300lf Hela cells, 2 × 10^4^ cells were seeded.

### Western Blotting

Cells were lysed in 2x Lamelli buffer (BioRad Cat. #1610737) with 5% β-Mercaptoethanol. Lysates were boiled and centrifuged prior to resolution on SDS-PAGE gels and transfer to a PVDF membrane. Membranes were probed with α-HiBiT (Promega 1:1000), α-NS6/7 (1:5,000), and α-GAPDH (Sigma, 1:1000).

### Anti-viral Screening

5 × 10^4^ BV2 cells per well were seeded in white-walled 96 well plates. At time of seeding, the cells were concurrently treated with indicated compound and either mock infected or infected with MNoV^CW3^-HiBiT at an MOI of 0.05. 24 hours post-infection, cells were subjected to HiBiT luciferase assay as described above. Viability of mock infected cells was determined using CellTiter-Glo (Promea) according to manufacturer’s instructions at 24 hours post infection.

## Data Availability

All relevant data are within the manuscript or in supporting information (S1 Table).

## Acknowledgements

We would like to thank Dustin Hancks, Craig Wilen, and all members of the Orchard Lab for helpful discussions. This work was supported by NIH grant 5R01DK133231 (R.C.O.).

## Author Contributions

M.C.O. designed the project, performed experiments and helped draft the paper. R.C.O. conceptualized the project, provided supervision, and helped write the paper. All authors read and edited the manuscript.

